# Host auxin machinery is important in mediating root architecture changes induced by *Pseudomonas fluorescens* GM30 and *Serendipita indica*

**DOI:** 10.1101/2020.08.17.251595

**Authors:** Anna Matthiadis, Poornima Sukumar, Alyssa DeLeon, Dale Pelletier, Jessy L. Labbé, Gerald A Tuskan, Udaya C Kalluri

## Abstract

Auxin is a key phytohormone that is integral to plant developmental processes including those underlying root initiation, elongation, and branching. Beneficial microbes have been shown to have an impact on root development, potentially mediated through auxin. In this study, we explore the role of host auxin signaling and transport components in mediating the root growth promoting effects of beneficial microbes. Towards this end, we undertook co-culture studies of *Arabidopsis thaliana* plants with microbes previously reported to promote lateral root proliferation and produce auxin. Two types of beneficial microbes were included in the present study; a plant growth promoting bacterial species of interest, *Pseudomonas fluorescens* GM30, and a well-studied plant growth promoting fungal species, *Serendipita indica (Piriformospora indica)*. Following co-culture, lateral root production was found restored in auxin transport inhibitor-treated plants, suggesting involvement of microbe and/or microbially-produced auxin in altering plant auxin levels. In order to clarify the role of host auxin signaling and transport pathways in mediating interactions with bacterial and fungal species, we employed a suite of auxin genetic mutants as hosts in co-culture screens. Our result show that the transport proteins PIN2, PIN3, and PIN7 and the signaling protein ARF19, are required for mediating root architecture effects by the bacterial and/or fungal species. Mutants corresponding to these proteins did not significantly respond to co-culture treatment and did not show increases in lateral root production and lateral root density. These results implicate the importance of host auxin signaling in both bacterial and fungal induced changes in root architecture and serve as a driver for future research on understanding the role of auxin-dependent and auxin-independent pathways in mediating plant-microbe interactions in economically important crop species.

## Introduction

Auxin is a key phytohormone that plays essential roles in above- and belowground plant developmental and physiological processes including embryogenesis, vascular patterning, seedling growth, apical dominance, tropic responses, and root growth and development (Enders and Strader, 2015; Taylor-Teeples et al. 2016). A modulation in concentration and location of auxin signal can dramatically influence plant growth and development. In belowground tissues, an internal genetically-mediated or external chemical increase in auxin concentration have been shown to inhibit root elongation and promote the development of lateral roots in a dose-dependent matter (Malamy 2005; Peret et al. 2009). The role of auxin has been further extended to its involvement in plant-microbe interactions in the root including nodulation and rhizobia symbiosis (Spaepen and Vanderleyden 2011; Grunewald et al. 2009) and non-nodule forming beneficial plant-microbe relationships (Sukumar et al. 2013; Boivin et al. 2016; Poupin et al. 2016). Specifically, microbes that can produce more auxin have been shown to have a greater beneficial impact on plant growth (Asghar et al. 2002; Khalid et al. 2004; Remans et al. 2007; Spaepen et al. 2014; Verbon and Liberman 2016). Recent studies have also demonstrated a link between microbe-induced lateral root formation and auxin or auxin-signaling pathways (Contreras-Cornejo et al. 2009; Zamioudis et al. 2013), but much remains to be characterized about the genetic implications and the underlying molecular mechanisms of these interactions, and whether specific molecular players are common to all microbe interactions or operate with species or strain specificity.

Our understanding of the role of auxin in plant-microbe interactions is limited and research efforts have largely included genetic and molecular investigations of pathogenic interactions, transcriptomics or proteomics changes associated with plant-microbe interactions, and a limited number of host genetic and mutant evaluations with beneficial microbes such as under nodulation (Grunewald et al. 2009; Zhao 2010; Sukumar et al. 2013; Contreras-Cornejo et al. 2015; Boivin et al. 2016; Poupin et al. 2016; French et al. 2018; Estenson et al. 2018). Auxin action can be regulated at the levels of biosynthesis, transport, and signal transduction (Lavy and Estelle, 2016). Given that increases in auxin synthesis may occur in microbes, outside of *in vivo* plant biosynthesis, investigation of transport and signal transduction within the plant can shed light on mechanisms of auxin-dependent/independent root system architecture alterations due to plant-microbe interactions.

Directional auxin transport is mediated by auxin influx proteins such as AUX1 (AUXIN RESISTANT 1) and LAX (LIKE AUXIN) and auxin efflux proteins including those belonging to PIN FORMED (PIN) and ATP BINDING CASSETTE B/MULTIDRUG RESISTANCE/P-GLYCOPROTEIN (ABCB/MDR/PGP) families (Teale et al. 2006; Kalluri et al. 2011; Enders and Strader, 2015). PIN2, PIN3, and PIN7 transport proteins have been implicated in lateral root formation in *Arabidopsis thaliana*. Specifically, *pin2*, *pin3*, and *pin2,3,7* mutants are known to have altered lateral root formation (Laskowski et al. 2008; Dubrovsky et al. 2009), PIN3 has been shown to be important for directing auxin to founder cells and promoting lateral root initiation (Marhavy et al. 2013), and *PIN3* and *PIN7* are known to be induced by IAA treatment (Lewis et al. 2011).

Auxin signaling components involve the auxin-binding receptor TRANPORT INHIBITOR RESPONSE 1 (TIR1) F-box proteins, which recognize AUX/IAA proteins as part of an E3 ligase receptor complex and ubiquitinate the proteins for 26S proteasome-dependent degradation. Removal of the repressor AUX/IAA protein from its binding partner protein, AUXIN RESPONSE FACTOR (ARF), triggers the ARF activity as a specific activator or repressor of auxin response gene transcription (Peret et al. 2009; Wang et al 2014; Roosjen et al. 2018). Several ARF isoforms have been implicated in regulating a variety auxin-dependent developmental processes in *Arabidopsis* (Roosjen et al. 2018). ARF7 and ARF19 have been reported to have partially redundant roles in the regulation of lateral root initiation through activation of LATERAL ORGAN BOUNDARIES-DOMAIN16 and 29 proteins (LBD16 and LBD29) (Okushima et al. 2005; Wilmoth et al. 2005; Okushima et al. 2007).

Past expression studies suggest functional roles for auxin transport and signaling proteins in root responses to plant-microbe interactions. *PIN2*, *PIN3*, and *ARF7* are upregulated in the colonization of *Arabidopsis* with the growth promoting bacteria *Burkholderia phytofirmans* PsJN (Sessitsch et al. 2005; Poupin et al. 2016), which has been recently reclassified under the genus name *Paraburkholderia* (Sawana et al. 2014). This study also employed a specific *Arabidopsis* auxin signaling genetic mutant, *iaa1/axr5-1*, to characterize co-culture effects (Poupin et al. 2016). Reduced lateral root induction was observed in *Arabidopsis* in co-culture with the ectomycorrhizal fungus *Laccaria bicolor* in *pin2* and *pin2, 3, 4, 7* mutants (Felten et al. 2009) and with the bacteria *Pseudomonas fluorescens* WCS417 in an *arf7arf19* double mutant (Zamioudis et al. 2013). Additional studies using more diverse beneficial microbial species, especially employing auxin genetic mutants, will help to elucidate the involvement of host auxin machinery.

Here we present new evidence supporting functional relevancy of host auxin transport and signaling components in mediating root architecture changes induced by non-nodule-forming, growth-promoting beneficial microbes. Specifically, we undertook co-culture screens in tissue culture using a suite of auxin genetic mutants followed by root architecture assessments. The targeted co-culture screens included two divergent species: *Serendipita indica (S. indica*; *Piriformospora indica)*, a fungal root endophyte, and *Pseudomonas fluorescens* strain GM30, a bacterial root endophyte. *S. indica* is a plant growth promoting microbe (PGPM) that has been widely shown to form mutualistic associations with and enhance growth in a variety of species including *Arabidopsis* (Verma et al. 1998; Peskan-Berghofer et al. 2004; Oelmüller et al. 2009). The bacterial isolate *P. fluorescens GM30* has been shown to alter plant development in *Arabidopsis* (Weston et al. 2012; Timm et al. 2015) and other plant species such as *Populus* (Henning et al. 2016). Both *S. indica* and *P. fluorescens* GM30 are capable of producing auxin and enhancing lateral root formation (Sirrenberg et al. 2007; Timm et al. 2015), however, the host genetic influence on these beneficial associations is unclear. Specifically, it is unclear if these root effects are dependent on auxin and host auxin molecular factors and whether these factors are broadly conserved between bacterial and fungal beneficial microbes.

## Materials and methods

### Plant and microbial materials and growth conditions, and plant phenotypic measurements

*Arabidopsis thaliana* seeds were surface sterilized and plated on ½ MS media with 10 g/L sucrose, 0.5 g/L MES, and 3 g/L Gelzan™ Gelrite® (Sigma-Aldrich) and incubated in a growth chamber under 16-h photoperiod at 22 °C. External auxin treatment included application of 10 μm 1-naphthaleneacetic acid (NAA) or 100 μm indole-3-butyric acid (IBA), and auxin transport inhibitor treatment included application of 100 μm 1-N-naphthylphthalamic acid (NPA) to plates containing 5-day old seedlings. Root measurements (total root length, number of lateral roots and primary root length) were recorded for 12-and 19- day- old seedlings (7- and 14- days following inoculation, respectively) using a WinRHIZO™ scanner (http://regent.qc.ca/assets/winrhizo_image_acquisition.html). Mutant seeds were obtained from ABRC (Arabidopsis Biological Resource Center) or from kindly donated by Dr Gloria Muday Laboratory and these include; *pin2* (*eir1-1* - CS8058) and *iaa14* (*slr-1* - CS25214) and from previously published research: *pin3pin7* (*pin3-5* - SALK_005544, *pin7-1* - SALK_048791), *arf7* (*nph4-1*), and *arf19* (*arf19-1* - CS24617). Control seedlings were generated from wild-type *Arabidopsis thaliana* (Col-0) seeds. *Pseudomonas fluorescens* strain GM30 and other bacterial strains were cultured overnight at 30 °C in R2A media (Teknova) and diluted to OD_600nm_ of 0.8 before inoculating plants by streaking*. Serendipita indica (Piriformospora indica*; belonging to *Hymenomycetes, Basidiomycota)* is a cultivable fungal endophyte that colonizes roots. *S.indica* has been studied for its multifunctional activities like plant growth promoter, immune modulator and biocontrol. *S. indica* was maintained at 30 °C in dark on modified Pachlewski medium P5 (Deveau et al., 2007). Three-weeks-old cultures were used and applied to seedling plates as plugs in ½ MS media with 10 g/L sucrose, 0.5 g/L MES, and 3 g/L Gelzan™ Gelrite® (Sigma-Aldrich) and growth chamber under 16-h photoperiod at 22 °C. Statistical significance was based on two-way ANOVA and p≤0.05. *Pseudomonas* strains include; *P. fluorescens* strains GM15, GM17, GM19, GM25, GM30, GM60, GM65, GM84 and GM89 from our previously reported collection (Jun et al. 2015; Timm et al. 2015), and *P. protegens* Pf-5/ATCC BAA-477, previously named as *Pseudomonas fluorescens* Pf-5 (Howell and Stipanovic, 1979), which was obtained from the American Type Culture Collection (Manassas, VA, U.S.A.) for this study.

## Results and discussion

### Application of exogenous auxin or beneficial microbes leads to lateral root proliferation

In order to define the baseline effect of exogenous auxin application or elevated auxin level on root architectural changes in wild-type *Arabidopsis thaliana* (Col-0) seedlings, auxin in the forms 1-naphthaleneacetic acid (NAA) and indole-3-butyric acid (IBA) were applied as agar streaks towards the bottom of the agar plate (Fig. 1a). The effect on root architecture was analyzed after seven days of treatment. Treatment with NAA and IBA resulted in enhanced lateral root emergence or proliferation. Increased lateral root production, with shorter lateral root length, was observed at seven days post co-culture with *S. indica* or GM30 relative to the uninoculated control (Supplemental Fig. 1), indicating that this effect could be auxin dependent. The auxin efflux inhibitor, 1-N-naphthylphthalamic acid (NPA), is known to arrest lateral root emergence (Petrasek et al. 2003). Hence, we examined the ability of these auxin-producing microorganisms to alleviate the auxin transport inhibitor-induced suppression of lateral root formation. NPA was applied locally at root-shoot junctions of 5-day old seedlings at the start of co-culture. Images were collected (Fig. 1b) and lateral root count was recorded (Fig. 1c) following seven days of treatment. While treatment with NPA resulted in no lateral root production in non-inoculated wild-type controls, addition of either *S. indica* or GM30 post-NPA treatment resulted in lateral root production (Fig. 1c). The restoration of lateral root formation with addition of *S. indica* or GM30 suggests that additional auxin signal, either in the form of microbial auxin and/or plant auxin, may be integral to pathways that modulate host root architecture in co-culture with these beneficial microbes.

**Fig 1.**
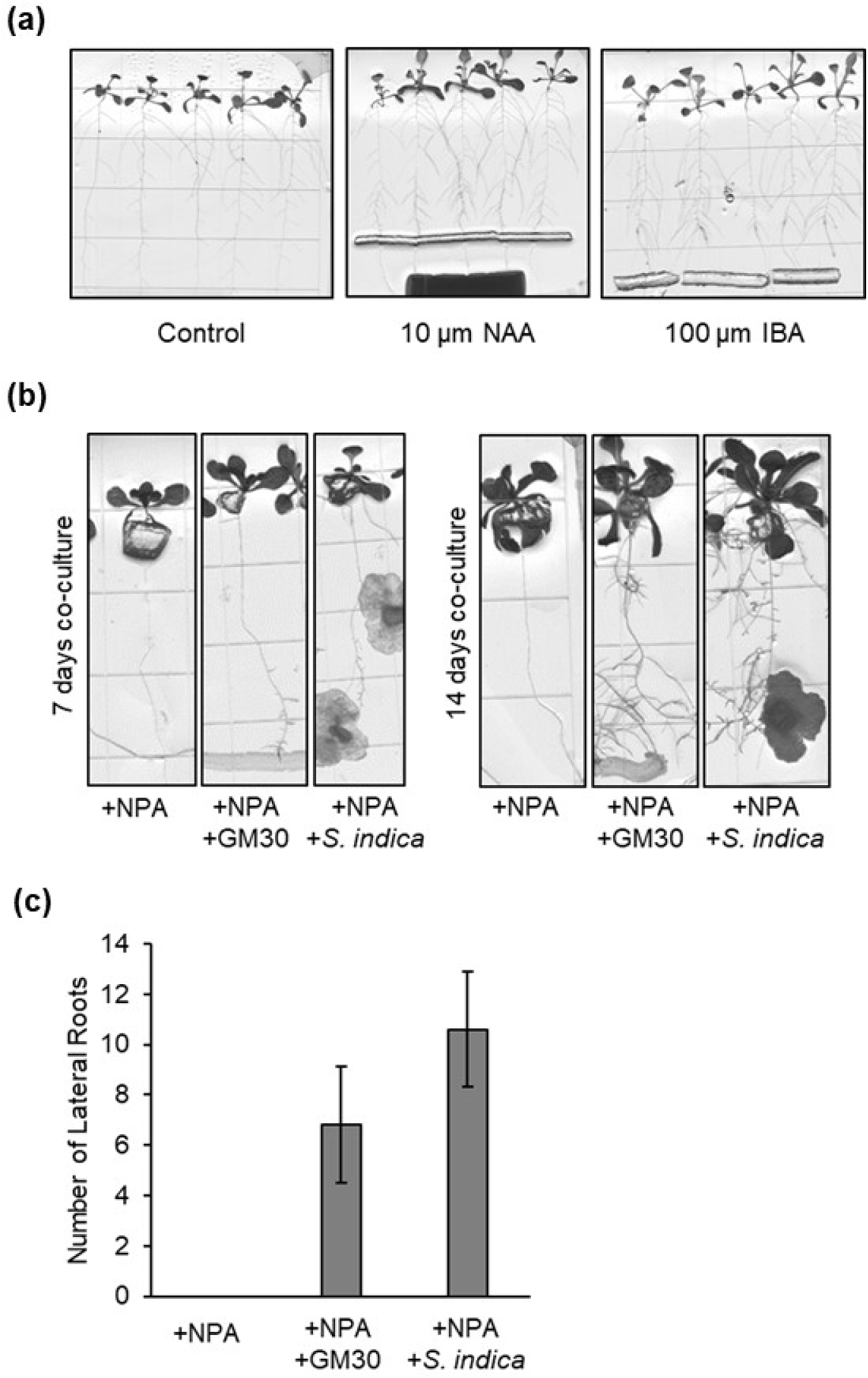
(a) Effect of exogenous auxin (NAA and IBA) application on root architecture. (b) Effect of auxin transport inhibitor NPA on lateral root number in Arabidopsis wild-type seedlings with or without co-culture (7 and 14 days). (c) Quantification of effect of NPA with or without co-culture on lateral root number (7 days).

### Auxin transport and signaling machinery is important in mediating microbe-induced root architecture changes

To gain an understanding of the core and differentiated components of host auxin pathways involved in bacterial- and/or fungal-mediated root architectural changes, we employed a genetic approach based on co-culture screens with known auxin genetic mutants. *Arabidopsis* mutants in auxin transport and signaling pathways were selected based on their previously reported involvement in formation or development of lateral or primary root growth. Changes in lateral root density induced by co-culture of *S. indica* and GM30 were examined and compared to wild-type plants (Fig. 2). The transport mutants displayed reduced response to co-culture-induced changes in root architecture. The *pin3pin7* double mutant displayed reduced responses to both microbes and *pin2* responded significantly to *S. indica* co-culture but not to GM30, relative to response in wild-type *Arabidopsis* (Fig. 2). The reduced *S. indica-* or *P. fluorescens*-induced host effects observed in transport pathway genetic mutants implicate the auxin transport machinery as integral to mediating growth promoting effects on the host via translation of changes in auxin levels into localized patterns of maxima and minima required for lateral root initiation. Our study also provides, to our knowledge, the first genetic evidence for auxin’s central role in mediating plant growth effects by *P. fluorescens* GM30.

**Fig 2.**
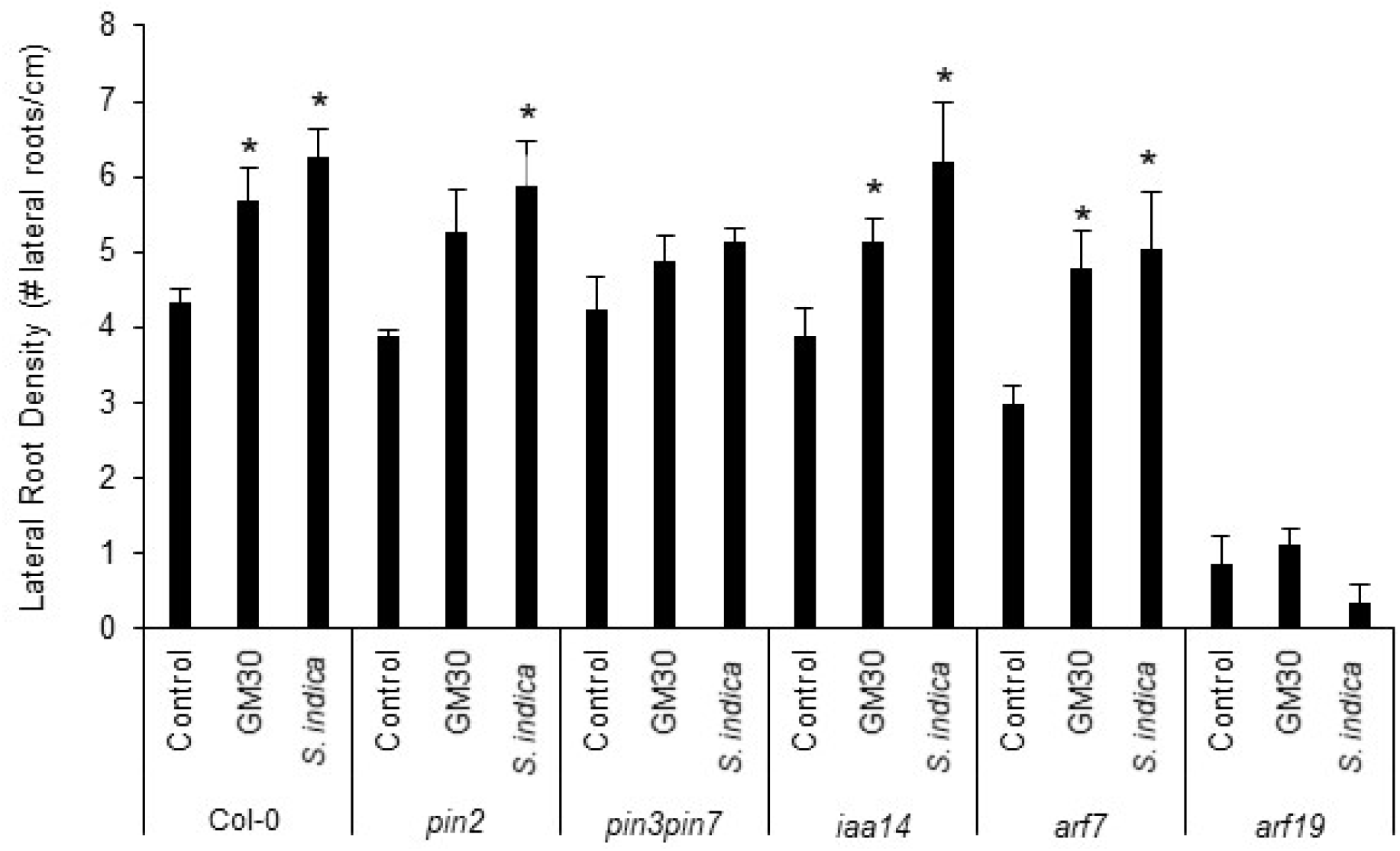
Effect of co-culture with *Pseudomonas flourescens* strain GM30 or *Serendipita indica* on lateral root density in *Arabidopsis* auxin transport and signaling mutant plants, relative to no microbe added control plants. Error bars indicate ±SE (n=5) and asterisks indicate significant difference from control (p≤0.05).

Among the auxin signaling mutants examined, *arf19* seedlings were found to be insensitive to co-culture, with either the bacterial or fungal symbiont (Fig. 2). The *arf19* mutant had a significantly lower lateral root density compared to wild-type even without microbial treatment, reflecting and confirming the well-studied role of ARF19 in lateral root development (Wilmoth et al. 2005). These results indicate that ARF19 may be critical not just in lateral root development but also in mediating root growth promoting effects of microbes. ARF7, whose function is reported to be partially redundant with ARF19 in lateral root development, also had a significantly lower lateral root density compared to wild-type under no microbe treatment yet was able to respond significantly to both microbial treatments (Fig. 2). *ARF19* has been reported to express more strongly relative to *ARF7* in roots (Okushima et al. 2005) supporting a plausible dominant role for ARF19 in microbe-mediated lateral root initiation.

The overall effects demonstrated are likely not specific to the *Pseudomonas* strain GM30 but may extend to other beneficial bacterial strains and species. Indeed, when testing a suite of additional auxin-producing *Pseudomonas* strains related to GM30 (Jun et al. 2015), we found a similar rescue in lateral root number in plants treated with NPA (Fig. 3), with all bacterial strains significantly increasing lateral root number in NPA treated plants and most strains restoring lateral root number to that of control (wild-type) plants not treated with NPA.

**Fig 3.**
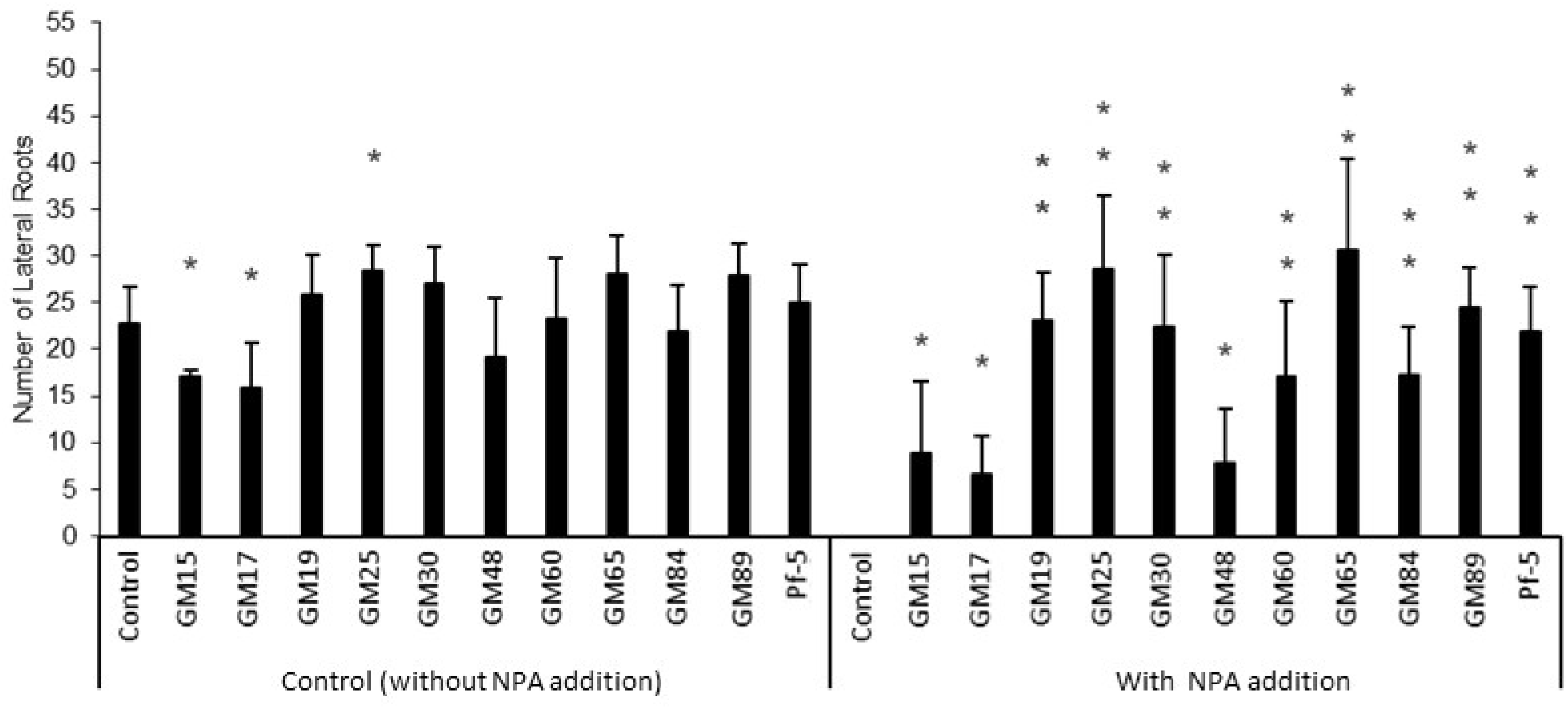
Effect of co-culture with various *Pseudomonas* strains on lateral root production in *Arabidopsis* wild-type seedlings, with or without addition of the auxin transport inhibitor NPA. *Pseudomonas* strains on x-axis include; *Pseudomonas flourescens* strains GM15, GM17, GM19, GM25, GM30, GM60, GM65, GM84 and GM89, and *Pseudomonas protegens* strain Pf-5 (Pf-5). Error bars indicate ±SE (n=5). One asterisk indicates significant difference from control plants without co-culture (p≤0.05). Two asterisks indicate NPA treated plants that are significantly different from NPA treated control plants without co-culture and not significantly different from control plants without NPA (i.e., restored to wild-type lateral root number) (p≤0.05).

## Conclusion

Our results implicate the importance of host auxin transport and signaling in microbe-induced changes in root architecture and set the stage for future research into the roles of auxin dependent and auxin independent pathways in mediating plant-microbe interactions in economically important crop species. Specifically, this study indicates a role for PIN3, PIN7, and ARF19 in mediating microbe-induced lateral root formation by both a bacterial and fungal species.

In the future, temporally-resolved experiments involving co-cultures with auxin mutant plants, application of external unlabeled or labeled auxins and auxin inhibitors, and examination of mutant phenotype rescue are needed to validate whether the observed changes are the consequence of adaptations in the plant system to support microbes or part of signal exchange between plant and microbes that leads to phenotypic changes. It is not known at this time whether the microbial auxin or additional biochemicals such as N-acyl-L-homoserine lactones (AHLs) and cyclodipeptides (CDPs) could be produced as a signal for individual microbes or for microbial communities and under what environmental and host-specific circumstances the functional outcomes change. Therefore, temporally-resolved molecular characterization and imaging would provide a clearer view of upstream and downstream events involved in the complex signal transduction process. Studies with a phylogenetically diverse range of bacterial strains to determine the range of specificity in the demonstrated auxin response is also needed. Finally, it is recognized that auxin action is mediated synergistically or antagonistically with other hormone signaling pathways including those involved in responses to external abiotic and biotic factors (Lewis et al. 2011; Boivin et al. 2016; Verbon and Liberman 2016), warranting a systems-wide view of plant-microbe interactions.

## Acknowledgements

We thank Prof. Gloria Muday and ABRC for provision of *Arabidopsis* mutant seeds used in the study. We thank Whitney McNutt and Daniel Smallwood for assistance with seedling cultures and data analysis, respectively. This research was sponsored by the Genomic Science Program, U.S. Department of Energy, Office of Science, Biological and Environmental Research as part of the Plant Microbe Interfaces Scientific Focus Area (http://pmi.ornl.gov). Oak Ridge National Laboratory is managed by UT-Battelle, LLC, for the U.S. Department of Energy under contract DE-AC05-00OR22725.

---

**Supplemental Fig 1.**
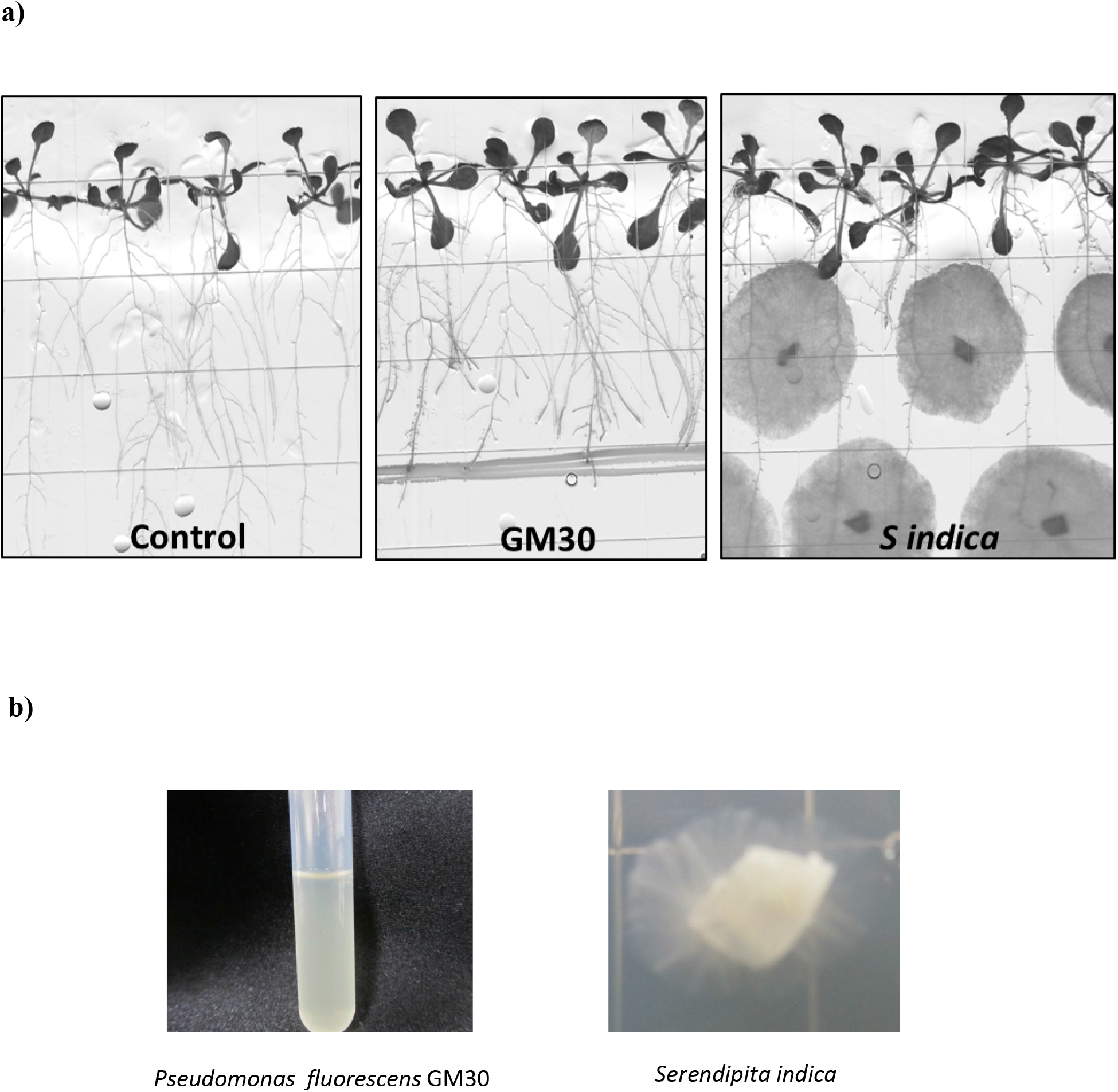
**a)** Effect on root architecture of wild-type *Arabidopsis* seedlings post- 7-day co-culture with **b)** *Pseudomonas fluorescens* GM30 bacterial or *Serendipita indica* fungal cultures.

